# ProteinVAE: Variational AutoEncoder for Translational Protein Design

**DOI:** 10.1101/2023.03.04.531110

**Authors:** Suyue Lyu, Shahin Sowlati-Hashjin, Michael Garton

## Abstract

There have recently been rapid advances in deep learning models for protein design. To demonstrate proof-of-concept, these advancements have focused on small proteins with lots of data for training. This means that they are often not suitable for generating proteins with the most potential for high clinical impact –due to the additional challenges of sparse data and large size many therapeutically relevant proteins have. One major application that fits this category is gene therapy delivery. Viral vectors such as Adenoviruses and AAVs are a common delivery vehicle for gene therapy. However, environmental exposure means that most people exhibit potent pre-existing immune responses to many serotypes. This response, primarily driven by neutralizing antibodies, also precludes repeated administration with the same serotype. Rare serotypes, serotypes targeting other species, and capsid engineering, have all been deployed in the service of reducing neutralization by pre-existing antibodies. However, progress has been very limited using conventional methods and a new approach is urgently needed. To address this, we developed a variational autoencoder that can generate synthetic viral vector serotypes without epitopes for pre-existing neutralizing antibodies. A compact generative computational model was constructed, with only 12.4 million parameters that could be efficiently trained on the limited natural sequences (e.g., 711 natural Adenovirus hexon sequences with average length of 938 amino acids). In contrast to the current state-of-the-art, the model was able to generate high-quality Adenovirus hexon sequences that were folded with high confidence by Alphafold2 to produce structures essentially identical to natural hexon structures. Molecular dynamics simulations confirmed that the structures are stable and protein–protein interfaces are intact. Local secondary structure and local mobility is also comparable with natural serotype behavior. Our model could be used to generate a broad range of synthetic adenovirus serotype sequences without epitopes for pre-existing neutralizing antibodies in the human population. It could be used more broadly to generate different types of viral vector, and any large, therapeutically valuable proteins, where available data is sparse.

## Introduction

As an increasing number of diseases have been associated with specific molecular pathogenesis, gene therapy is gaining more attention as it provides targeted treatment from the source. Efficient and targeted gene delivery is an important step that can greatly influence the treatment effect (Bulcha et al., 2021). Different delivery vehicles have been designed to increase efficiency and specificity. Naturally, viral vectors are an attractive class of tool for gene delivery, since viruses are highly evolved to efficiently gain access to the host. Viral vectors have been engineered to exclude the replication and toxicity related coding regions (Thomas et al., 2003) for safety consideration. There are several commonly used viral vectors, including lentivirus, retrovirus, herpes simplex virus 1, adenovirus (Ad), and adeno-associated virus (AAV). Adenovirus vectors have several advantages, including: 1) delivered DNA exists episomally, avoiding potential oncogenesis and other issues that can arise from genomic integration; 2) large packaging capacity of 36kb for delivery gene circuits or treating diseases involving pathogenic mutation of multiple genes; 3) highly efficient transduction (60-80%) of most dividing and quiescent cells(Bouvet et al., 1998; Chillon et al., 1999; Stevenson et al., 1997); 4) technology is readily available for scalable vector production(Bulcha et al., 2021; Thomas et al., 2003). For these reasons, 50% of all ongoing gene therapy clinical trials are using Adenovirus-based vectors (Bulcha et al., 2021). However, with their natural coexistence with human, wild-type virus vectors often trigger complex immunological reactions that greatly hinders treatment efficiency, and thus pre-existing immunity is their main limitation (Lee et al., 2017; Thomas et al., 2003). Pre-existing neutralizing antibody against the most used adenovirus strain (AdV5) was found in over 30% of tested sera samples from Europe, Asia, and the United States (Vogels et al., 2003). Another study revealed that anti-AdV2 neutralizing activity existed in over 40% of tested children’s sera samples, and 74% of children’s sera samples contained neutralizing antibodies for at least one serotype(D’ambrosio et al., 1982). In addition, transient expression is typically observed for genes delivered episomally, and therefore, multiple rounds of adenovirus-mediated therapy will likely be required (Lee et al., 2017). To prevent immunity associated with repeated administration, different serotypes are needed for each round. As the major capsid protein, hexons were identified as the primary target of neutralizing antibodies (Sumida et al., 2005), Chemical or biological modifications to the adenovirus capsid hexons can assist in evasion of the serotype-specific neutralizing antibodies (Seregin and Amalfitano, 2009). However, success thus far has been incremental and limited. A more aggressive design approach is needed to generate Adenovirus vectors with no epitopes for neutralizing antibodies. Changing the entire solvent exposed surface such that all potential neutralizing antibody epitopes are removed, involves introducing a large number of mutations (∼ 452 mutations). Introducing this many mutations randomly or using rational structure based design cannot be done without catastrophic loss of protein function (Ogden et al., 2019; Sarkisyan et al., 2016). We hypothesized that machine learning could be used to generate dramatically different hexon proteins without impacting essential folding and function.

Deep learning models have been recently developed for de novo protein design (Castro et al., 2022; Ding et al., 2019; Hawkins-Hooker et al., 2021; Nijkamp et al., 2022; Ogden et al., 2019; Repecka et al., 2021; Riesselman et al., 2018; Sevgen et al., 2023). However, these models involve a large number of parameters and thus require a large amount of data for training. For example, ProteinGAN, the first Generative Adversarial Networks model designed for protein sequence generation, comprised 60 million trainable parameters and was trained on 16,706 unique malate dehydrogenase (MDH) sequences (Repecka et al., 2021). As there are only 88 serotypes of human adenovirus identified so far (Dhingra et al., 2019), the number of available unique hexon sequences are limited to 711 unique full-length sequences from the UniprotKB database. Hexon sequences are 938 amino acids long on average, which suggests there is likely to be more inter-residue dependency occurring at longer distances. No previous work has reported generating sequences of comparable length. This, combined with the small dataset, meant that a small but expressive model would be required –– that can be trained efficiently. To solve this, a pre-trained protein language model for amino acid-level embedding was used, allowing transfer of knowledge learned by the pretrained model on a large protein database. A Variational Autoencoder (VAE) (Kingma and Welling, 2013) framework was used to obtain an informative and structured latent space, and thereby convert the discrete protein sequence space to a continuous space for ease of sampling and other manipulation. A special bottleneck attention module (Montero et al., 2021) was used in the encoder to map the high-quality amino-acid-level embedding–––generated by the pretrained model–––to the latent space. A non-autoregressive deconvolution-based decoder was designed for sequence reconstruction from the latent variable. Our final model was able to generate high quality, structurally stable hexon sequences with only 12.4 million parameters.

## Results

To collect training data, all hexon proteins in the UniprotKB database that are annotated to be full-length were extracted. Since the goal is to train a model for generating full-length hexons with all domains, sequences that are likely to have incomplete domains were removed with a length cutoff of 800 amino acids. To accommodate ease of downstream application, only sequences with standard amino acids are kept in the dataset. The resultant dataset includes 711 hexon sequences with length ranging from 893 to 992 amino acids. To illustrate the generative capacity, a recently published large transformer-based language model ProtGPT2 (Ferruz et al., 2022) was fine-tuned on the same hexon dataset. Another common approach is to train models for designing sequences with a fixed-backbone as input. The state-of-the-art fixed-backbone design method requires prohibitively large computational resources (Verkuil et al., 2022). Instead, ESM-InverseFolding (ESM-IF1), a competitive model, was included for comparison as a representative model for this approach. ESM-IF1 was not fine-tuned due to the lack of structural data for most sequences in the training dataset.

### Generative VAE Model for Small Protein Families

Given the limited amount of data available for certain protein families, the aim was to develop a model that can be effectively trained on a small dataset. Inspired by(Khandelwal et al., 2019; Montero et al., 2021), a pre-trained protein language model was incorporated in the encoder, as illustrated in Figure 1. A convolutional neural network was used to extract a feature vector from the hidden amino-acid-level representation produced by the pretrained protein language model (Supplementary Figure 1a). Using the CNN-extracted feature vector as the query in the bottleneck attention module, global-level information was integrated to obtain a refined protein-level representation. Instead of training from scratch on the limited hexon data, the parameters in the pretrained model were kept fixed. This permitted use of the high-quality amino-acid-level embeddings and more efficient training on a small dataset. ProtBert was chosen as the pre-trained protein language model, because it is the best performing model that was trained on sequences with length ranges (2048 amino acids) that cover hexons.

**Figure 1.**
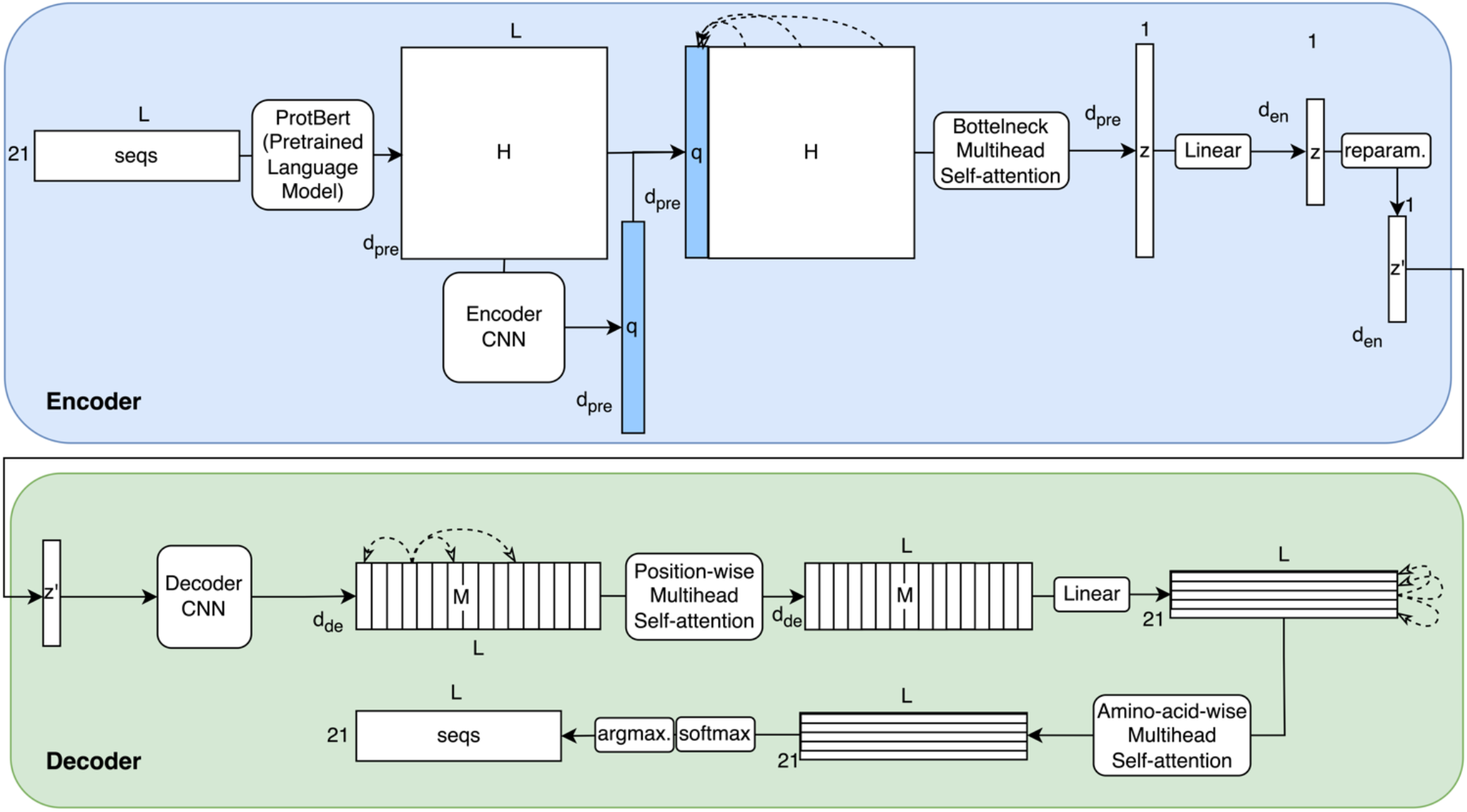
ProteinVAE architecture: Encoder: Attention mechanism with fixed query vector ([CLS] vector) is used to extract bottleneck representation *z* from pretrained language model hidden representation *H*. Decoder: Deconvolutional networks are used for upsampling. Attention modules are applied on both dimensions.

For the decoder, a non-autoregressive processing method was used for reasons mentioned above. Inspired by the Hybrid VAE paper (Semeniuta et al., 2017), deconvolutional layers (Supplementary Figure 1b) were designed to upsample the bottleneck representation, protein-level representation, to amino-acid-level representation *M* of size *L* × *d*_*de*_, where *L* means sequence length, and *d*_*de*_ means decoder hidden dimension. Specifically, a convolution layer with kernel size of 1 × 1 separated deconvolutional layers with a bigger kernel (Iandola et al., 2014). This resulted in a reduced number of parameters needed. Next, a position-wise multi-head attention mechanism was used to capture the dependencies between amino acid usage at different positions, which allowed effective modeling of the long-distance interaction. A linear layer was used to convert the hidden representation *M* to the logits matrix. Another multi-head attention module is designed to adjust for amino acid preference in different viruses (Bahir et al., 2009), and therefore, it is done across different amino-acid channels. Lastly, some initial results of the model generated more helix sequences than strand (Supplementary Figure 2). Combined with the consensus that strand proteins are harder to design, a reweighted cross-entropy loss that assigned higher penalty to the strand positions, predicted by SPOT-1D (Hanson et al., 2019), was used in the final model.

**Figure 2.**
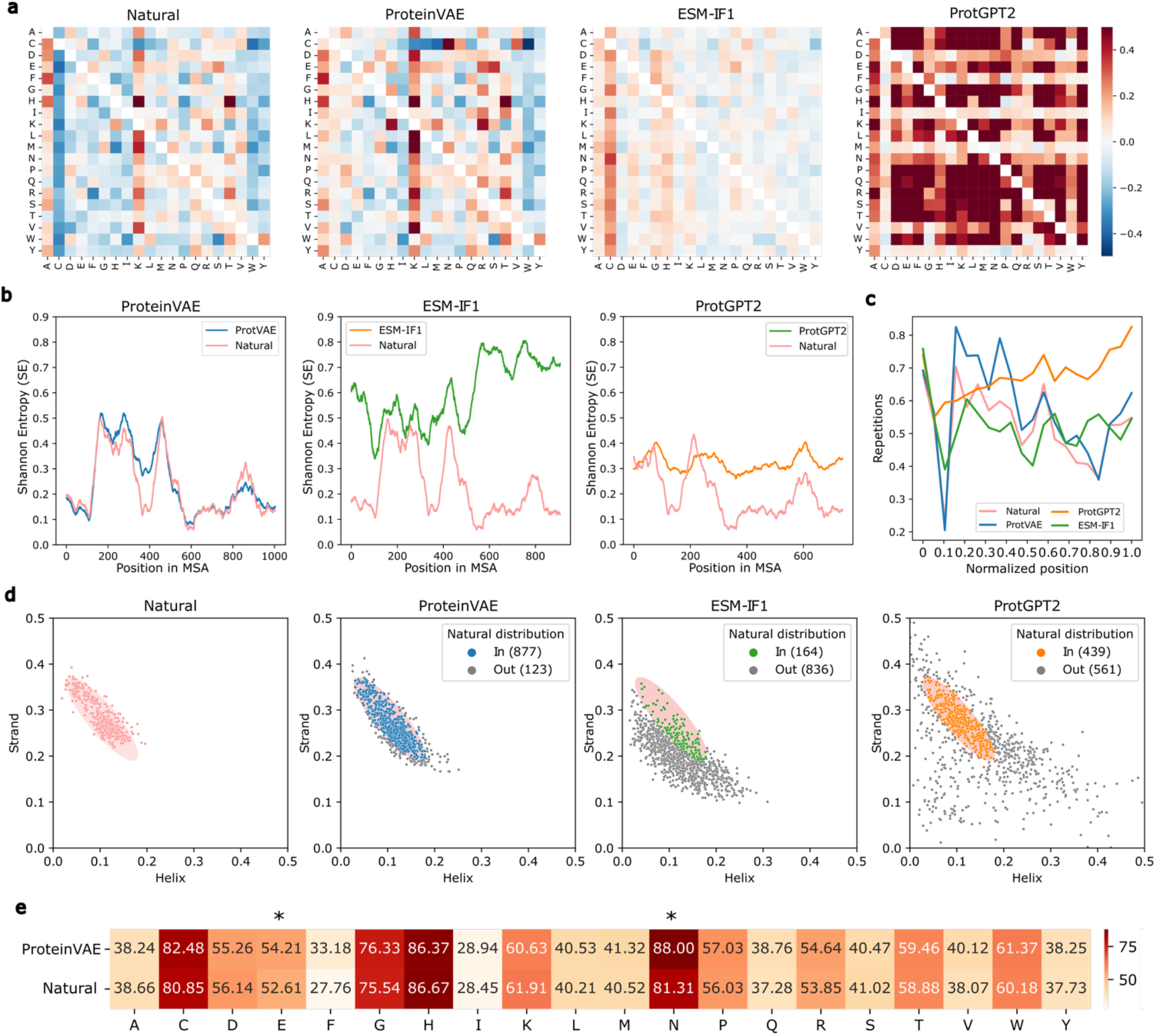
**(a)** Amino-acid pair association scores for all ProtGPT2 generated, ProteinVAE generated, and natural sequences. Negative values (blue) indicate shorter distances compared to random shuffled sequences. **(b)** Shannon-entropy for natural hexons and sequences generated by all models in MSA columns with above 20% occupancy in each datasets. Higher value reflects higher sequence variability across samples. **(c)** Average repetition in sliding windows of length 5. Higher value means more repeated amino acids are present in a window. X-axis shows the normalized position of the left end of the window. **(d)** Helix and strand ratio in natural and generated hexons. Pink shade in all plots shows the area in the bi-variate normal distribution fitted on natural samples (*α* = 0.05). In generated sequence plots, gray points represent outliers, while colored points are sequences considered within the natural distribution. (**e)** SASA for all amino acids in ProteinVAE generated and natural proteins. Star indicates amino acids with significantly different (*α* = 0.05) SASA values between two groups.

### ProteinVAE Generates Sequences Similar to Natural Hexons

To evaluate sequence generation, 1,500 samples were generated by sampling a Gaussian distribution with mean of 0 and standard deviation of 4 from the latent space and decoded by the ProteinVAE decoder. Top 1,000 sequences with highest likelihood were used for analysis. For comparison, 1,000 sequences were also generated using ESM-IF1 and fine-tuned ProtGPT2, respectively. For the ESM-IF1 model, 10,000 sequences were generated using 100 Alphafold2 predicted hexon structures as template. Top 1000 sequences with highest log likelihood were selected for comparison. Even after fine-tuning, ProtGPT2 could not consistently generate protein sequences within the length range of the training data. 42,550 sequences were generated with fine-tuned ProtGPT2, and only ∼ 3% of all sampled sequences were within the length range. To meaningfully compare the generated sequences, ProtGPT2-generated sequences within the length range were ranked according to perplexity, and top 1,000 sequences with lowest perplexity were selected.

To assess ProteinVAE’s capacity of learning the distribution of natural hexon sequences, several metrics were computed comparing both local and whole-sequence level patterns. As an initial evaluation, local amino acid positional patterns were examined. Amino-acid pair association score (Santoni et al., 2016) (based on minimal proximity function) was calculated for all possible combinations of amino acids in natural sequences and sequences generated by 3 models (Figure 2a). Similarity of the pair association score between ProteinVAE generated sequences and natural sequences indicated that ProteinVAE has learned the local amino acid patterns of natural hexons. In comparison, almost all amino acid pairs occur at a closer distance than randomly shuffled sequences in fine-tuned ProtGPT2 generated sequences, which is substantially different from the patterns observed in the natural sequences. Fewer associated pairs are detected in ESM-IF1 generated sequences, as indicated by the overall low absolute value, which also differs from the natural pattern.

Next, the generated sequences were assessed for preservation of the natural evolutionary profile, namely, sequence conservation and variability in specific regions. Shannon entropies (SE) were computed for all valid positions in the multiple sequence alignment (MSA) of natural sequences and sequences generated with different models, as shown in Figure 2b. SE of ProteinVAE generated sequences presented peaks and valleys at similar locations to the natural sequences (Pearson’s *r* = 0.90), which indicated that ProteinVAE has learned the underlying sequence distribution. Fewer columns were preserved for SE calculation in the ProtGPT2 and ESM-IF1 generated sequences, because more invalid columns with more than 80% gaps existed in the MSA. Different sequence variability patterns were also present in the ESM-IF1 (Pearson’s *r* = 0.15) and fine-tuned ProtGPT2 (Pearson’s *r* = 0.50) generated sequences. This suggested that the homology between natural hexons and sequences generated by ProtGPT2 and ESM-IF1 is more distant.

Sequence length is one of the characteristics of hexons that added difficulties to its modeling. As reported before, generating long (∼1000s tokens) and coherent texts in a specific small domain is challenging even for fine-tuned large language models like GPT2 (Holtzman et al., 2019; Tan et al., 2020), and generated texts typically suffer from degenerate repetition. To evaluate if the generated sequences can avoid the degenerate repetition artifacts while capturing certain local repetitive patterns observed in natural sequences (Jorda et al., 2010), number of repeated amino acids was calculated in a fixed-length window sliding across all possible positions in each sequence (Figure 2c). Regardless of the window size used (Supplementary Figure 3), ProteinVAE samples closely follow the repetitiveness trend as the natural (Pearson’s *r* = 0.94), while ESM-IF1 generated sequences showed a slightly different repetition pattern (Pearson’s *r* = 0.52). The repetition increases as the generation progresses in ProtGPT2 samples, introducing more repetitive artifacts towards the end of the sequence (Pearson’s *r* = 0.12).

**Figure 3.**
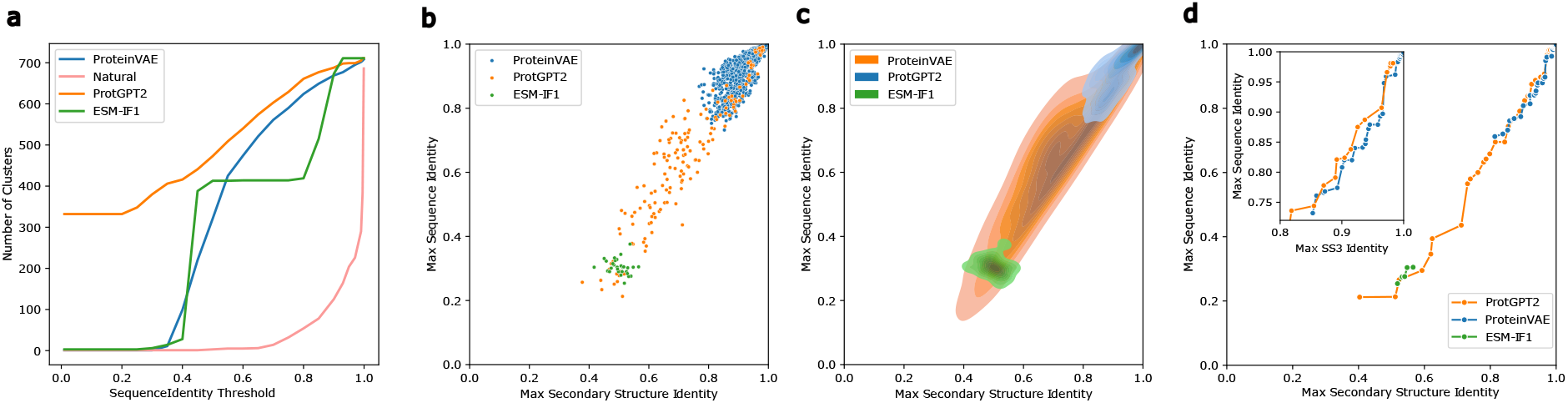
**(a)**Number of clusters at different identity thresholds. **(b)** Sequence diversity plotted against secondary structure similarity. X-axis is the maximum percentage identity of 3-state secondary structure on all aligned pairs of generated and natural sequences. Y-axis is the maximum sequence identity on all aligned pairs. Sequences closer to the bottom-right corner are ideal, as they are structurally similar to natural protein but more novel in sequence. **(c)** Density plot for sequence diversity and secondary structure similarity. **(d)** Pareto frontiers: The optimal sequences designed by two models are highlighted along respective frontier.

After validating the model’s capacity in generating sequences resembling natural hexon proteins with sequence similarity analysis, secondary structure composition of the generated sequences was analyzed. First, the Q3 secondary structure was predicted for all natural and generated sequences with SPOT-1D (Hanson et al., 2019). Since the ratio of coil is dependent on the ratio of strand and helix (sum of all three ratios is 1), only the strand and helix percentage were analyzed. As shown in Figure 4a, the strand and helix ratio are correlated (Pearson correlation *r* = − 0.86) in natural hexons. Similar trend existed in the ProteinVAE samples (Pearson correlation *r* = − 0.86), but not in the ProtGPT2 samples (Pearson correlation *r* = − 0.60) nor ESM-IF1 samples (Pearson correlation *r* = − 0.73) (Figure 2d). To further analyze secondary structure profile, a bivariate normal distribution was fitted on the natural set, and out-of-distribution samples were identified in the generated sequences (*α* = 0.05). More samples generated by the ProteinVAE model (877 sequences) share similar secondary structure composition with natural hexons, compared to ESM-IF1 samples (164 sequences) and ProtGPT2 samples (439 sequences) (Figure 2d).

**Figure 4.**
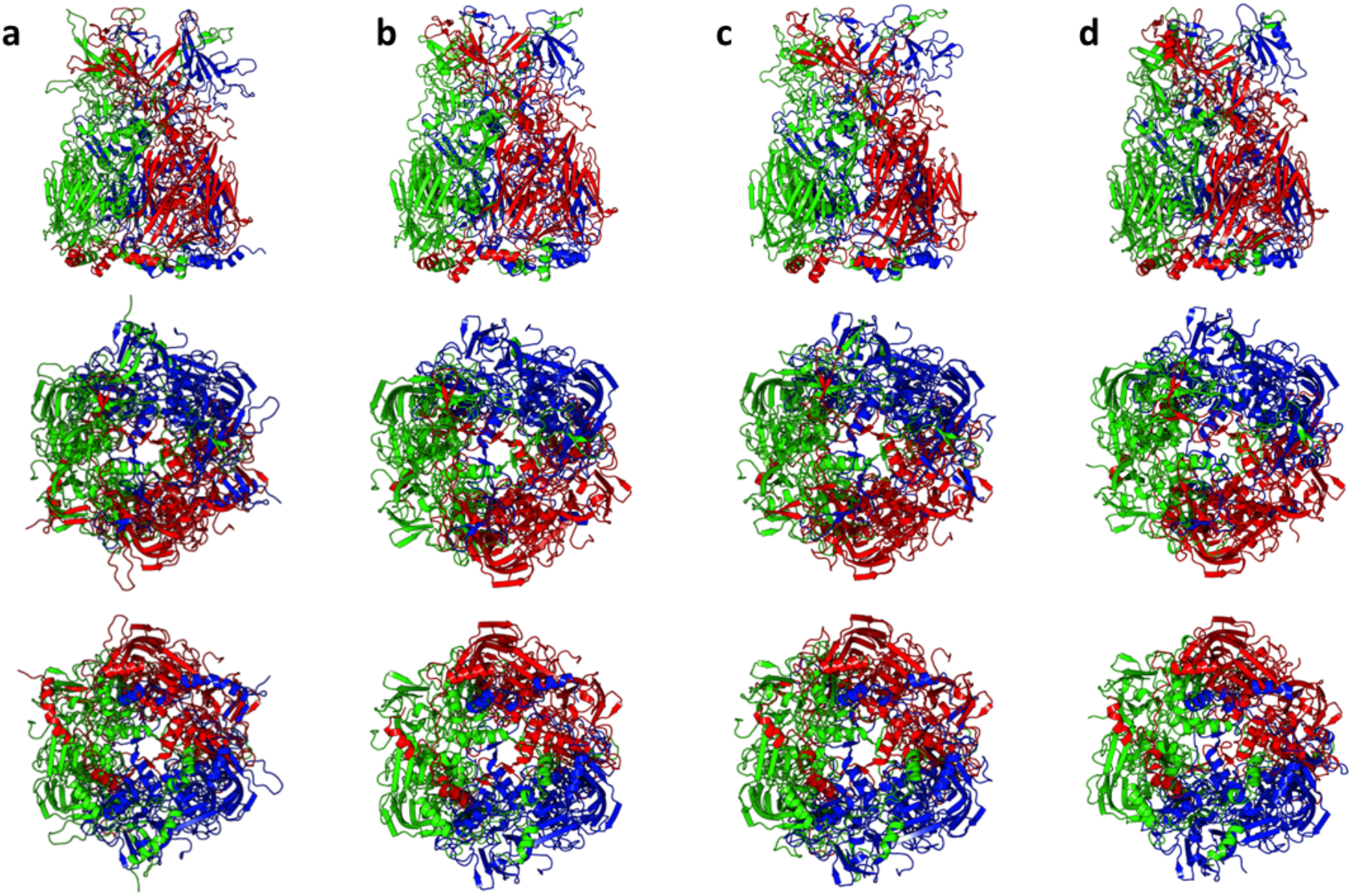
MD representative structures of **(a)** WT and ProteinVAE generated sequences with **(b)** 91.5%, **(c)** 85.6%, **(d)** 75.4% sequence identity with respect to their respective closest natural sequence. Side, top, and bottom views of all structures were shown in the first, second, and third row, respectively.

To further analyze the structure profile, solvent accessible surface area (SASA) profile was computed for all 20 amino acids in 100 randomly selected sequences from ProteinVAE samples and natural hexons, respectively (Figure 2e). SASA was calculated from Alphafold2 (Jumper et al., 2021) predicted structures. SASA profile for 18 amino acids are similar in natural and ProteinVAE generated sequences. 2 amino acids (E, N) have significantly different SASA values, but their surface exposureness is retained. This comparison supports that ProteinVAE generated sequences are structurally similar to natural hexons. Only ProteinVAE generated sequences were analyzed due to limited computing resources.

### ProteinVAE Generates Diverse Sequences

To visualize sequence diversity, an equal number of natural sequences and generated sequences by all models were clustered at different thresholds. Although, as discussed in the previous section, ProteinVAE generated sequences closely resembling sequence patterns found in natural hexon populations, they consistently had more clusters and higher diversity than natural sequences at all thresholds (Figure 3a). In ProtGPT2 generated samples, there exists a large number of clusters even at extremely low identity threshold. This is likely the result of ProtGPT2 inserting sequence fragments from the vast Uniref50 database that it was pre-trained on (Ferruz et al., 2022).

Furthermore, since the goal is to generate functional hexons that are biologically relevant, the ability of the model to diversify the sequences while keeping high structure resemblance towards natural hexon protein is critical. As shown in Figure 2d, more samples generated by ProteinVAE had similar secondary structure composition to natural protein. Additionally, to assess sequence diversity meaningfully, sequences with less than 80% target and query coverage when aligned to its closest natural sequence were removed, as this increases the chances of the generated sequences having most domains required for functionality and decreases the possibility of involving non-hexon domains that could introduce unwanted effects in future gene therapy. All ProteinVAE samples satisfy this sequence-level constraint, as the ProteinVAE model has only been trained on hexon sequences. In contrast, only 163 samples are left in the ProtGPT2 group, which further supports our hypothesis that ProtGPT2 incorporated non-hexon fragments in generated sequences. While the absolute sequence identity of ProteinVAE samples only covers the higher end of the range seen in ProtGPT2 samples, all 877 analyzed ProteinVAE samples have high structural similarity towards natural sequences (Figure 3b-c). This is likely because only the top 1000 sequences with the highest average positional likelihood were selected from the 1,500 sequences generated by the ProteinVAE model, which ensures high sequence quality at the cost of lower diversity. There were only 30 sequences in ESM-IF1 generated samples that satisfied the screening conditions, and all remaining samples had lower secondary structure similarity towards natural hexons. To directly evaluate sequence diversity against structural similarity, we plotted the Pareto frontier of the all generated samples, respectively in Figure 3c. In the comparable range, ProteinVAE produced samples more diverse without disruption of structural profile.

### Molecular Dynamics Simulations

Molecular dynamics simulations were used to assess structural stability of generated hexon trimers. To obtain the range of conformational flexibility in natural hexons, we performed clustering on the natural sequences at 90% sequence identity and collected the representative sequence from all clusters with more than 10 sequences (13 clusters). To evenly sample the sequence space, 50,000 sequences were sampled with the mean of all latent vectors from each cluster and a standard deviation of 3. Sequences that are more repetitive than the natural sequences from the respective cluster were filtered. From the remaining sequences, sequences of different identities (60 - 92%) towards the closest natural protein were selected for molecular simulation. For comparison, sequences were randomly selected from ProtGPT2 generated sequences that are above 80% in both query and target coverage when aligned with the closest natural sequence. 3 ProtGPT2 generated sequences were randomly selected in each 10% identity range from 70% - 100% (9 sequences in total).

3D structures for all sequences were predicted with Alphafold2. Hexon forms homotrimers in the adenovirus capsid (Figure 4). The three monomers form a quasi C3v point group with extensive interactions between the monomers. On average, the 13 representative natural hexon monomers consist of ∼ 18% helix and 28% β-strand, with turns and coils making up the remaining 54% of the structure (Drew and Janes, 2020). A comparison between the representative structure of a select natural hexon (A4ZKL6) and three ProteinVAE designed hexons (with 91.5%, 85.6%, and 75.4% identity to the respective closest natural hexon) reveals that all three designed structures maintained the overall shape and symmetry of the natural counterpart (Figure 4).

All predicted structures were subject to classical molecular dynamics simulation for 100 ns. Root-mean-square deviation (RMSD) of all natural hexons and all sequences generated by ProteinVAE and ProtGPT2 are within a stable range (natural hexon: 1.14 − 4.53*Å*, ProteinVAE samples: 1.21 − 6.58*Å*, ProtGPT2 samples: 1.23 − 8.86*Å*) (Supplementary Figure 4). Root-mean-square fluctuation (RMSF) of the structures were analyzed to identify changes in local structural flexibility (Figure 5). As can be seen, ProtGPT2 samples introduced mutations that significantly increased sequence flexibility in sequence regions that were relatively rigid in natural sequences, whereas mutations introduced in ProteinVAE samples do not suffer from this artifact. In addition, ProtGPT2 samples inserted long fragments that are not homologous to natural hexons, and these fragments have high flexibility that could decrease structural stability (Supplementary Figure 3). Mutations in ProteinVAE samples (Figure 5 c) are more likely to occur in regions that are exposed (Figure 5 d) and more likely to be tolerant towards mutations.

**Figure 5.**
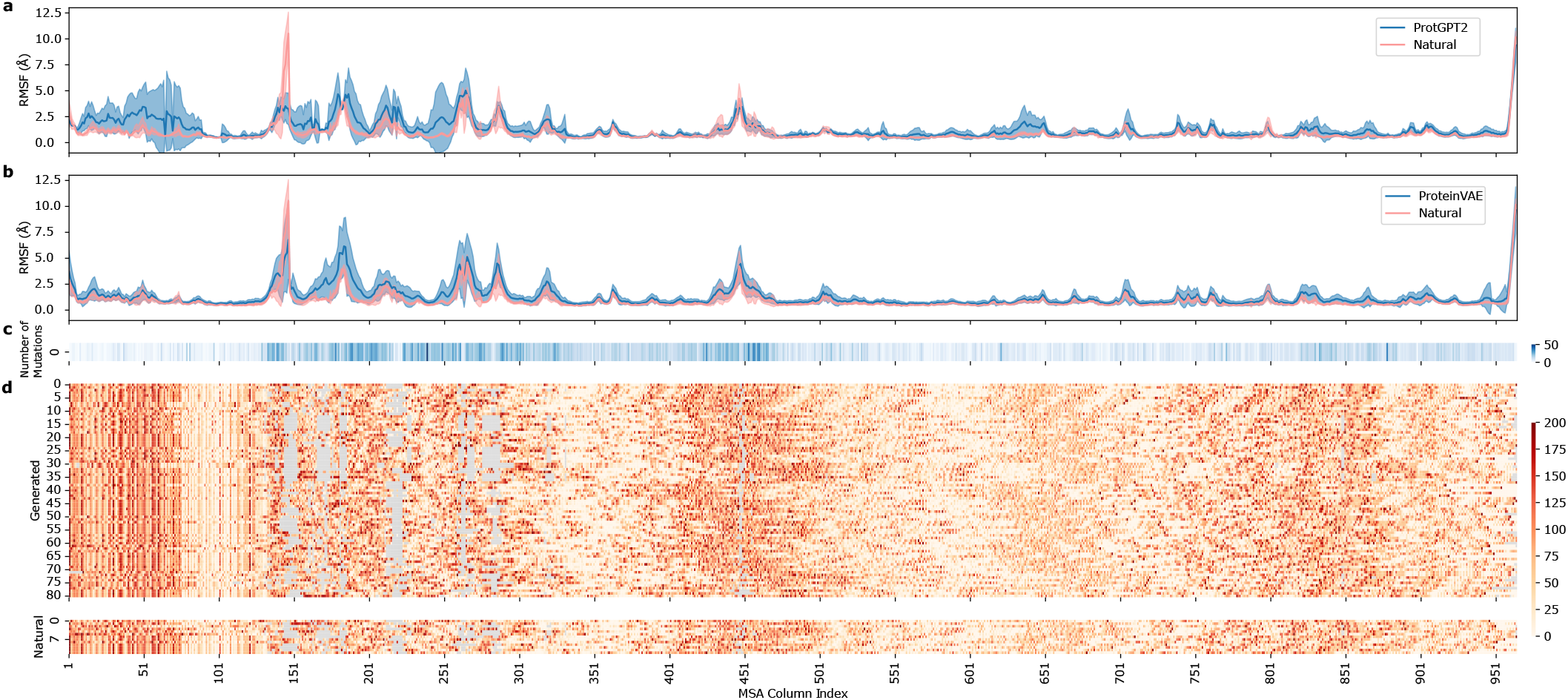
**(a)** RMSF for all wild-type cluster representative sequences and stable ProtGPT2 generated sequences. **(b)** RMSF for all wild-type cluster representative sequences and stable ProteinVAE generated sequences. **(c)** Heatmap of positions where mutations were introduced in stable ProteinVAE generated sequences compared to their closest natural sequence, respectively. **(d)** Heatmap of solvent accessible area across all positions in each natural sequence and stable ProteinVAE sample.

### Novel Synthetic Human Adenovirus Serotype

To distinguish human adenovirus serotypes from generated sequences, a simple logistic regression classifier was trained from the encoder embeddings of all training data. The ROC AUC of the trained classifier is 0.97 (Figure 6a). Next, sequences generated from each cluster were encoded and classified (Figure 6b). Percentage of generated sequences classified as human adenovirus hexon correlated with that of natural sequences in each cluster (pearson’s r = 0.81). Next, the sequences that are classified as human adenovirus hexon were aligned with all-natural human adenovirus hexon with known serotype. Amino acid divergence in loop 1 and loop 2 were calculated for each pair of sequences (Figure 6c). Generated sequences diverged more than 4.2% in loop 1 and more than 1.2% in loop 2 from any known serotypes, which suggested that they are likely hexons from new human adenovirus serotypes(Madisch et al., 2005).

**Figure 6.**
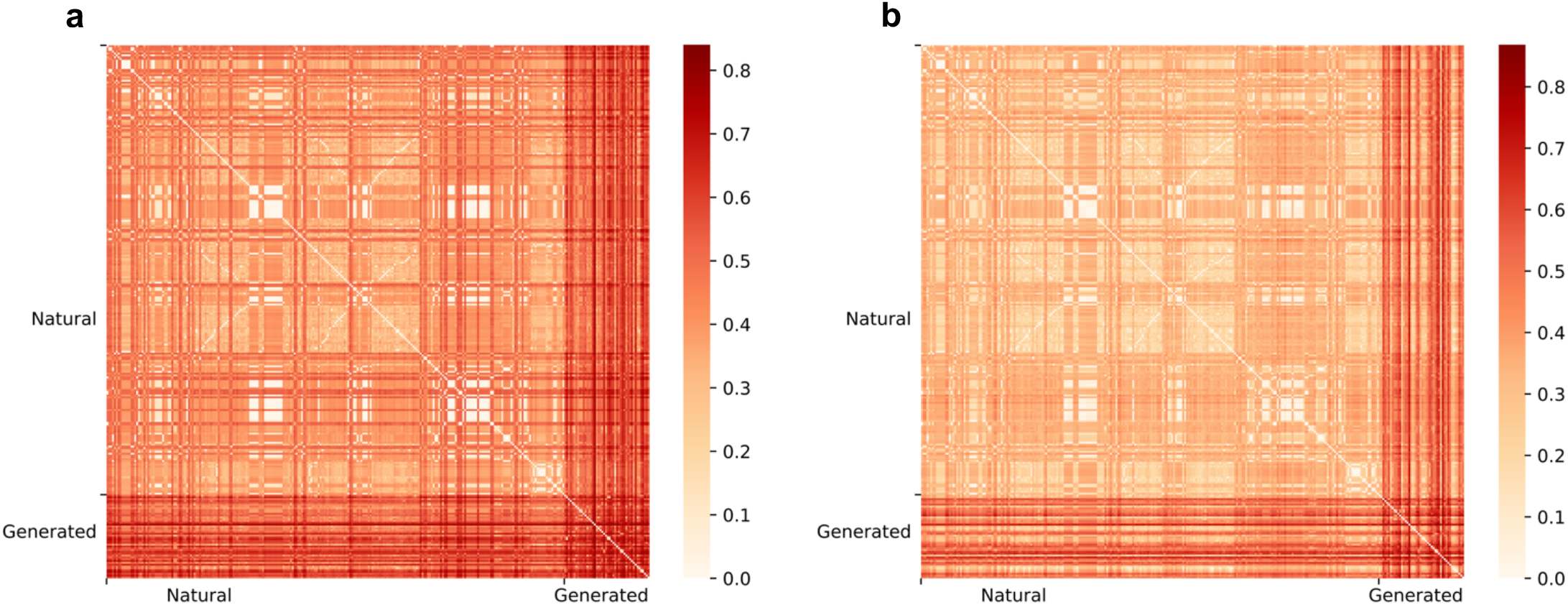
**(a - b)**. Pairwise amino acid divergence for the loop1 (column 129-390 in MSA) and loop2 regions (column 447-540 in MSA) in hexon, respectively. Darker shade means more difference between the compared sequences.

### ProteinVAE Latent Space Allows Interpolation between Natural Sequences

One benefit of using the VAE-based model is the ease of sampling provided by the structured VAE latent space. To validate that evolutionary relationships and sequence similarities have been captured in the latent vectors, the 10 largest clusters (at 90% sequence identity threshold) were plotted in dimension-reduced hidden space (obtained with Principal Component Analysis) (Figure 7a). Multiple clusters can be found in the hidden space distinctly separated. ProteinVAE hidden space also appears around the mean of 0 with no obvious hole.

**Figure 7.**
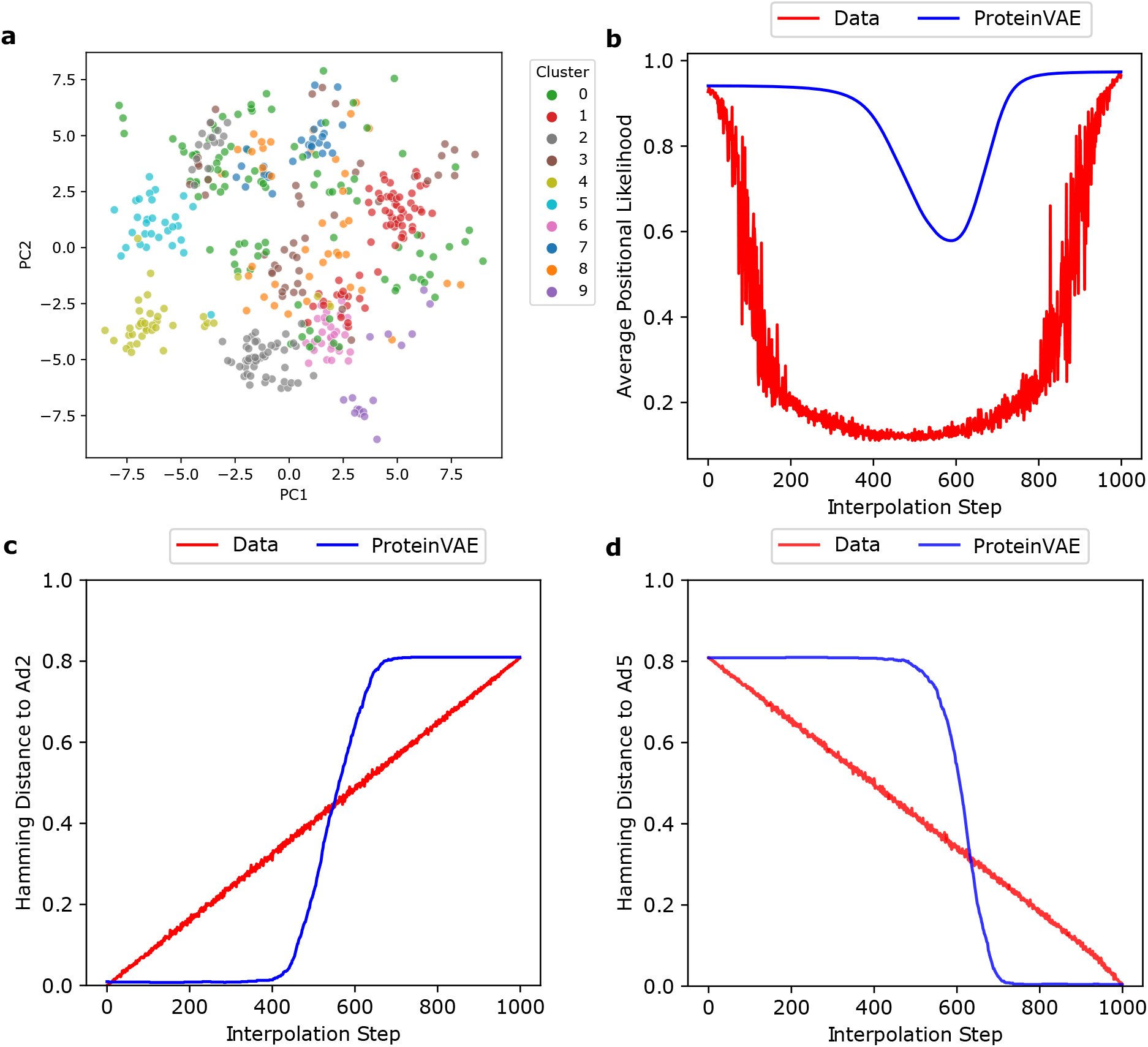
**(a)** Latent space clustering of sequences from 10 largest clusters at 90‥ identity. **(b)** Average positional likelihood for latent space interpolated sequences and direct interpolated sequences. **(c - d)** Hamming distance to ad2 and ad5 hexon, respectively.

Next, interpolation was done between hexons from two adenovirus serotypes, AdV2 and AdV5, which are experimentally-tested to be interchangeable (Youil et al., 2002). As mentioned in the introduction, AdV5 is the most common adenovirus vector, which means creating mutants to avoid its pre-existing neutralizing antibodies would be more impactful than creating mutants for a random natural hexon. Hexon exchange between certain serotypes could result in unsuccessful virus assembly (Youil et al., 2002). By interpolating between exchangeable hexons, there are increased chances of forming viable viruses with resulting hexon variants and the rest of the proteins from the AdV5 genome. 1,000 vectors were linear interpolated between AdV5 and AdV2 hexon hidden vectors in ProteinVAE latent space, since this is a common approach to utilize VAE structure latent space (Roberts et al., 2018; Wang et al., 2022). The interpolated vectors were decoded to sequences. As a control, data interpolation approach mentioned in (Roberts et al., 2018) was implemented to sample between AdV5 and AdV2 hexon sequences directly. Both the control and latent interpolation achieved monotonic changes in hamming distance (Figure 7c-d). However, ProteinVAE latent interpolation allows for generating of natural-resembling sequences, as indicated by higher average positional likelihood (Figure 7b). This example showcased another method of applying the ProteinVAE model to aid rapid sequence design based on known biological constraints.

### Ablation Study

The goal of this study is to design a model that is intrinsically easy to sample where only latent variables are required during generation. This would also allow convenient sequence manipulation in latent space. Thus, instead of passing amino-acid-level embedding to the decoder, any encoder-decoder information flow other than the bottleneck vector was eliminated. Then, the generation from the bottleneck vector can be viewed as an upsampling process. Similarly, when provided with the same input vector at each time-step, traditional LSTM-based (Sundermeyer et al., 2012) sequence generation can also be viewed as an upsampling process. To test the hypothesis that non-autoregressive generation is more suitable in the case of protein design, deconvolution-based upsampling was also compared with the traditional LSTM method. Aftering tuning the number of layers, processing direction, and hidden dimension, the best performing LSTM-based model (14.0M parameters) only achieved a reconstruction accuracy of 0.18 on the validation set, while the deconvolution-based (12.4M parameters) model achieved 0.86. As an alternative to using LSTM as an upsampling method, training LSTM with additional per-time-step token information in a teacher forcing manner was experimented, and this led to detrimental overfitting. No generation with the LSTM-based model was attempted, given the extremely low reconstruction accuracy.

To deepen our understanding of each component’s contribution to the generation capacity, an ablation study was conducted by leaving out non-standard elements of the network one at a time. Reconstruction and generation performance was summarized for each version of the ablated model in table 1. The full model performs best overall. The encoder CNN module improved the reconstruction accuracy, which is likely due to the addition of local features that reflects detailed differences among sequences. The bottleneck attention module has more effect on improving generation, which is likely tied to its ability to produce a better protein level representation that complies with the sampling process (Montero et al., 2021). Surprisingly, the amino acid attention improved generation in terms of secondary structure ratio and sequence variability profile, but not the amino acid usage. In addition, the secondary structure reweighting on the cross-entropy loss only improved amino acid usage and sequence variability profile slightly. This could be related to the error in the secondary structure prediction.

**Table 1.** Ablation study, each settings are repeated with 3 different random seeds. Encoder CNN: CNN feature vector extractor in encoder; Bottleneck attn.: Autobot attention mechanism (Montero et al., 2021); Amino-acid-wise attn.: Attention mechanism across different amino-acid channel; Secondary-structure-weighted
CE Loss: cross entropy reweighted to assigned 1.2x penalty to the strand positions; Test F1: reconstruction F1 score on test set; JSD (Amino acid usage): Jesen-Shannon distance between amino acid usage frequency;JSD (Secondary Structure): Jesen-Shannon distance between secondary structure ratio; MSA Entropy Correlation: Pearson correlation between MSA entropies in valid columns of generated and natural sequences.

|  | Encoder<br>CNN | Bottleneck<br>attn. | Amino-acid-wise<br>attn. | Secondary-structure-weighted<br>CE Loss | Test F1 | JSD (Amino<br>acid usage) | JSD (Secondary<br>Structure) | MSA Entropy<br>Correlation |
| --- | --- | --- | --- | --- | --- | --- | --- | --- |
| ProtGPT2 (fine-tuned) | / | / | / | / | / | 0.040 | 0.073 | 0.521 |
| Final | + | + | + | + | 0.861 | 0.022 | 0.015 | 0.911 |
| w/o encoder CNN | - | + | + | + | 0.836 | 0.023 | 0.015 | 0.878 |
| w/o bottleneck attn. | + | - | + | + | 0.868 | 0.039 | 0.038 | 0.500 |
| w/o aa attn. | + | + | - | + | 0.847 | 0.013 | 0.024 | 0.669 |
| w/o ss weighted CE | + | + | + | - | 0.858 | 0.023 | 0.012 | 0.902 |
| Base | - | + | - | - | 0.790 | 0.015 | 0.044 | 0.436 |

## Discussion

ProteinVAE can be used to learn the intrinsic relationships of a long protein sequence from a limited number of samples. Computational evaluation between ProteinVAE and a fine-tuned large language model, ProtGPT2, revealed that ProteinVAE can more reliably capture population level statistical characteristics of natural hexons. Specifically, ProteinVAE generated sequences that resembled natural sequences on both sequence level and structural level. In addition, ProteinVAE generated sequences are more diverse than natural sequences, capable of forming more clusters at the same identity threshold. ProteinVAE has a structured latent space which can be used for sequence interpolation between biologically validated interchangeable hexons. Lastly, using only 711 training samples, ProteinVAE was able to generate diverse sequences that are tested to be stable in molecular simulations, with the most novel sequence containing 291 amino acids different from its closest natural sequence with 39.62% viral surface area changed.

Considerable efforts have been made toward computationally expanding known protein families with novel sequences. In conventional bioinformatics, hidden Markov models (HMMs) are used to capture pairwise coupling within the MSA of a protein family, and as an extension, it can be used to generate new sequences. However, previous research has shown inconsistent levels of success with this type of model (Repecka et al., 2021; Russ et al., 2020), mostly due to its inability to learn higher-order relationships that exist in natural protein families. As an alternative, modern deep learning models, including Generative Adversarial Networks (GAN) (Goodfellow et al., 2020; Repecka et al., 2021), VAEs(Hawkins-Hooker et al., 2021; Riesselman et al., 2018; Sevgen et al., 2023; Sinai et al., 2017), and large generative protein language models (Ferruz et al., 2022; Madani et al., 2023; Nijkamp et al., 2022), have been implemented to learn the complex constraints in biological sequence design. However, previous methods have mostly focused on shorter protein sequences with a large number of members from the same family. The difficulties with the task of generating diverse hexon sequences were further demonstrated with the unsatisfactory performance of a fixed-backbone design model (ESM-IF1), and a recently-published large language model (ProtGPT2) fine-tuned on hexon dataset. Although the state-of-the-art fixed-backbone-design method showed more promising results on smaller proteins, these models require significantly higher GPU memory for generating long sequences, as the inter-residue distance information requires quadruple amount of memory for processing as the length increases. For instance, 24GB GPU memory is needed to generate 1 hexon sequence using ESM-IF1. Only 21GB GPU memory is needed to generate 1000 hexon sequences in parallel for ProteinVAE. For sequences of even longer length, generation might not be feasible for ESM-IF1, as the memory requirement for generating a single sequence might exceed typical GPU memory. It is also possible to improve the ProtGPT2 generation, but it would take significantly more computational resources to search for the optimal generation settings. Instead, ProteinVAE distilled knowledge from a pre-trained protein language model and leveraged it to facilitate efficient learning of the complex sequence patterns from limited data. Moreover, the ProteinVAE model was able to generate 1000 sequences in less than 1 minute, while the generation of 1000 lower-quality sequences took ∼13 hr for the ProtGPT2 model. Lastly, the generated sequences analyzed above were selected with an emphasis on sequence quality, which limited the range of diversity to some extent. In the future, less stringent selection criteria could be used to obtain more diverse sequences.

Concurrent works, ProT-VAE (Sevgen et al., 2023) and ReLSO (Castro et al., 2022), both involved autoencoder and the use of language model, but 1) neither presented results on designing protein at the same length range as hexons, 2) both models were trained for a different objective of exploring fitness landscape and generating functionally improved sequences, 3) both used larger labeled datasets (ProT-VAE: 6447 and 20,000 sequences, ReLSO: 10^10^, 20^4^, and 51,175 sequences). Briefly, ProT-VAE incorporated a generic CNN network for compressing and decompressing pre-trained language model (PLM) hidden states, and the protein-family-specific VAE part was trained to further reduce the protein-level representation to a single vector and reconstruct the hidden states given the vector. The VAE part was trained with maximum mean discrepancy loss instead of KL-divergence.

ProT-VAE is built on a ProtT5 model that was pre-trained on sequences with a maximum length of 512 amino acids. Although, T5 model was previously demonstrated to be able to extrapolate up to another 600 tokens when trained on 512 tokens in natural language processing tasks (Press et al., 2021), it is unclear if ProT-VAE can be used to design longer proteins like hexon. Since the ProT-VAE model has not been released, we simulated the reconstruction F1 in ProT-VAE model by introducing a small Gaussian noise to the encoder output before decoding. Since the ProtT5 model (48 M parameters) used in ProT-VAE was not publicly available, another larger ProtT5 (3 B parameters) was used. As a comparison, 5000 phenylalanine hydroxylase (PAH) sequences were found via psiBLAST with a human PAH variant (2PAH) as described in ProT-VAE. Owing to the high computational resources required by running the large T5 model, an equal size PAH dataset was randomly selected from the psiBLAST results for comparison. When a small Gaussian noise (variance < 0.0625) was injected to the encoder output, the simulated reconstruction performance on the PAH dataset remained at a high level (F1 = 0.81).

When tested on the hexon dataset, the T5 model can faithfully decode all hexon sequences in the training dataset (F1 = 1) without noise. However, when a smaller level of Gaussian noise (variance < 0.04) was added, significantly worsened F1-score (0.19) was observed on the hexon dataset (Supplementary Figure 7). Because of the significantly low performance with noise, it is likely that the ProT-VAE will not be able to generate high-quality hexon sequences using a ProtT5 pre-trained on sequences less than 512 amino acids.

Due to the model design in ProT-VAE, exchanging the PLM would not only require retraining the VAE part on family-specific sequences, but also retraining the generic CNN network on a large protein database which is computationally prohibitive. In comparison, accommodating another PLM in ProteinVAE only requires training the VAE component on sequences from a specific protein family. It is also unlikely that a pre-trained PLM decoder can be used directly to reconstruct hexon, because this small family is likely underrepresented in the pre-training database. Moreover, in (Sevgen et al., 2023), the diversity of sequences was only measured against one natural sequence. This could lead to overestimation of sequence novelty, because the generated sequences could be very different from the compared single natural sequences while sharing high identity with other natural sequences.

The ReLSO model was designed with an autoencoder, instead of a VAE architecture, and fitness information is jointly trained to be encoded in the latent space. It was trained on both positive and negative samples with a specially designed interpolation loss. The language model was not pre-trained in ReLSO. ReLSO is likely to be less effective in designing hexons due to limited data.

The capacity of ProteinVAE to learn the complex protein sequence distribution from limited samples could potentially be applied in a variety of different sequence design problems. Traditionally, exploration of the protein fitness landscape has been done with mutagenesis libraries (Sarkisyan et al., 2016). This approach is inherently inefficient as only neighboring regions can be explored. In the future, another model could be trained to map the ProteinVAE latent space to the protein fitness landscape, and apply the ProteinVAE model to conditionally generate sequences with functional improvement(Ding et al., 2019). Such computational exploration may facilitate exploration of distant regions of the fitness landscape where significant functional enhancement might be. Pre-training on larger relevant protein datasets could also potentially increase sequence diversity.

## Method

### Dataset

Hexon protein is the major capsid protein in adenovirus with a length spanning from 893 to 992 amino acids, with an average length of 938 amino acids. To increase the chances of generating complete sequences covering all domains, only full-length Hexon proteins annotated in the UniprotKB database were collected. These sequences were then filtered for those shorter than 800 amino acids for quality purposes, and for ease of downstream application, sequences with non-standard amino acids (U, J, Z, O, B, X) were removed. In total, 711 hexon sequences were collected as training data. The same training/validation/test set splits ratio 7/2/1 were used for all models. The same random seed is used for each replicate group respectively.

### Variational Autoencoder

Variational Autoencoder (VAE) (Kingma and Welling, 2013) is composed of an encoder and a decoder. The encoder *qϕ*,(*Z*|*x*) a neural network parameterized by *ϕ*, maps the input data samples *x* into a latent variable *z*, assumed to follow a Gaussian distribution as its prior. The decoder *pϕ* (*x*|*z)*, another neural network parameterized by *θ*, reconstructs the sample *x* from the latent variable *z*. VAE is trained by maximizing ELBO, where

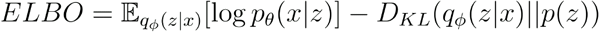

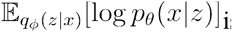 is the expected conditional log-likelihood. *p*(*z*) is the prior Gaussian distribution, *D*_*KL*_(*q*_*φ*_(*z*|*x*)||*p*(*z*)) is the KL-divergence. The details are described in (Kingma and Welling, 2013).

In common text generation tasks, it has been demonstrated before that when KL-divergence decreases too much, the generated samples are likely to suffer from low diversity (Bowman et al., 2015). To prevent KL-vanishing and to allow effective manipulation of the impact of KL-divergence, a non-linear proportional-integral-derivative (PID) controller was implemented to automatically tune the weight of KL-divergence in the VAE objectives throughout training. The KL-divergence weight *β*(*t*) is calculated through a feedback control defined as:

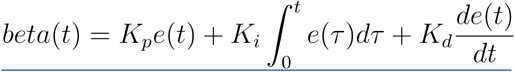

*e*(*t*) is the error between actual and expected value at time *t. K*_*p*_, *K*_*i*_ and *K*_*d*_ are the coefficients for the proportional, integral, and derivative term, respectively. Details can be found in (Shao et al., 2020).

### Bottleneck Encoder

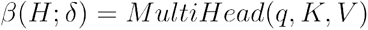

where the keys *K* (size *T × d*) and values *V* (size *T × d*) are transformed from the output hidden representations of the pretrained language model *H* (*T × d*). The parameter *δ* includes the weights for transformation of the query, keys, and values. During training, the pretrained language model stays frozen, only the parameter is *δ* learned.

Restricted by the length of hexon, only limited pretrained language models are available. ProtBert (420M parameters) (Elnaggar et al., 2020) was chosen as the pretrained protein language model, as 1) it is trained with a length limit of 2048 amino acids, 2) it achieved better results on downstream tasks than other models trained on long protein sequences. In brief, the ProtBert version used in ProteinVAE contains 30 layers, and it was trained for 300k steps on sequences shorter than 512 amino acids, then for an additional 100k steps on sequences with a maximum length of 2k (Elnaggar et al., 2020).

### Non-autoregressive Decoder

As mentioned in the introduction, the hypothesis was that the protein sequence can be better modeled with a non-autoregressive processing approach. Thus, inspired by the HybridVAE model (Semeniuta et al., 2017), deconvolution networks were used to perform upsampling. The deconvolutional network increases the spatial size of the input, while decreasing the number of hidden dimensions. Specifically, the deconvolutional networks consist of 8 UpBlocks. In each UpBlock, a 1 × 1 convolution layer transforms the input to have a lower number of channels which reduces number of parameters needed in the next layer; a 3 × 3 deconvolutional layer upsamples the low-channel input. To maintain gradient, the output of each previous UpBlock was concatenated as the input for the next block. Unlike HybridVAE, deconvolutional networks output *M* was not passed to a recurrent neural network (RNN). Instead, a multi-head attention module was used to capture both short and long range relationships. Next, the output was converted to logits using a linear layer. Lastly, another amino-acid wise attention module was added to capture the amino acid usage preference among different viruses (Bahir et al., 2009). As a comparison, classic autoregressive processing was tested by replacing the deconvolutional networks with a multi layer long short-term memory (LSTM) RNN (Sundermeyer et al., 2012). Both single direction and bidirectional LSTM models are tested. LSTM hidden sizes are divided by 2, when testing bidirectional LSTM. The hidden dimensions of the bottleneck representation *z* and the upsampled decoder hidden representation *M* were kept the same.

### ProteinVAE Training

ProteinVAE model was trained on the hexon dataset using negative ELBO loss. KL-divergence was dynamically weighted using a PID controller with expected KL-divergence of 0.5, *K*_*p*_ of 0.01,*K*_*i*_ of 0.0001, and *K*_*d*_ of 0.001. Strand position weighted to have 1.2x cross-entropy loss. ProteinVAE model was optimized using Adam optimizer (Kingma and Ba, 2014) with a learning rate of 0.0005 and weight decay of 0.0001. Drop out rate of 0.3 was used. A one-cycle learning rate scheduler (Smith and Topin, 2019) was used with a total step of 8000, percentage of the cycle (in number of steps) spent increasing the learning rate was set to 0.4, and the initial learning rate was set to 1/20 of peak learning rate. Decoder and encoder latent size are set to 128. In each of the 8 decoder UpBlock, upsampling was done after input was transformed to 16 channels. Encoder bottleneck attention has 4 heads. Decoder position-wise attention has 2 heads, and the decoder amino-acid-wise attention also has 2 heads. ProteinVAE model was trained on a NVIDIA V100 GPU with 32GB memory.

### ProteinVAE Sequence Generation

#### Generate from cluster for molecular dynamic analysis

For each of the top 13 clusters (90% identity, size>10), 50000 sequences were generated with the mean of respective cluster and standard deviation of 3. Within each cluster, sequences more repetitive than the most repetitive natural sequence in that cluster were filtered out. For the rest of the sequences, each one is aligned against the whole natural hexon dataset to get the percentage identity towards the closest natural protein. A binwidth of 2% was used to separate sequences of different novelty. For the 17 bins with percentage identity from 60%-92% (some bins are empty), the sequence with the highest average position probability is selected for AF2 structure prediction. Structures with an average pLDDT score of higher than 85% were selected for molecular dynamic analysis.

#### All other samples are generated following this procedure

To increase sequence diversity, vectors were samples from a normal distribution with mean of 0 and standard deviation of 4, and decoded them to new sequences. To maintain high sequence quality, sequences were ranked according to average positional likelihood, and selected the top 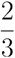 sequences for downstream analysis. Average positional likelihood (APL) was calculated as 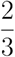 sequences for downstream analysis. Average positional likelihood (APL) was calculated as:

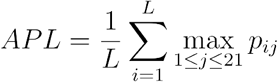

Where *p*_*ij*_ is the predicted probability of amino acid *j* (including a special token “-” representing a gap in sequence) at the *i*th position. *L* is the maximum length of training sequences. *p*_*ij*_ is the predicted probability of amino *j* acid (including a special token “-” representing a gap in sequence) at the *i* th position. *L* is the maximum length of training sequences.

### Alphafold2 structure prediction

Due to limited computing resources, all structures were predicted with the AF2 reduced database. After folding a small number of natural hexons, it was observed that model_1 predictions ranked the first for all sequences. To save computing time, only model_1 was used for all predictions reported here. All predictions were run on NVIDIA A100 GPUs with 48GB memory.

### Multiple Sequence Alignment Entropy

Clustal Omega (Sievers and Higgins, 2018) was used to calculate multiple sequence alignment (MSA) for the entire natural hexon dataset mixed with an equal number of generated sequences. Columns with more than 80% gaps in either natural or generated dataset were removed. Shannon entropy within each column was calculated as:

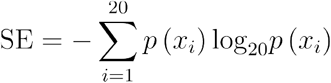

Where *p*(*x*_*i*_) is the frequency of amino acid in *i* each column. Pearson correlation was calculated between valid entropy values of natural and generated sequences. *p*(*x*_*i*_) is the frequency of amino acid *i* in each column. Pearson correlation was calculated between valid entropy values of natural and generated sequences.

### Association Measure for Amino-acid Pairs

For any pair of amino acid *a* and *b*, the minimal proximity score and the pair association metrics were calculated as described in (Santoni et al., 2016). Distance between each occurrence of *a* at position *x*_*i*_ with its nearest occurrence of *b* at position *y*_*i*_ was computed, and then averaged across all occurrences of *a*:*a* and *b*, the minimal proximity score and the pair association metrics were calculated as described in (Santoni et al., 2016). Distance between each occurrence *a* of at position *x*_*i*_ with its nearest occurrence of *b* at position *y*_*i*_ was computed, and then averaged across all occurrences of *a*:

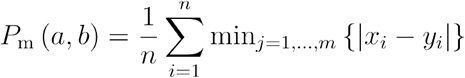

To remove dependency on the number of occurrences for different amino acids, the position of *a* was fixed and *b* was randomly shuffled. Mean (*p*_*m*_(*a*,Rand(*b*))) and standard deviation 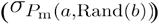 of the randomly shuffled sequences (where position of *b* is shuffled, but position of *a* is fixed) were calculated, and the minimal proximity score was normalized to obtain the association score as described in (Santoni et al., 2016).

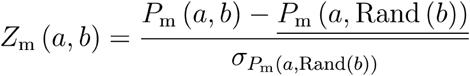

For the score shown in Figure 2, averaged association scores were plotted for each group of sequences. A null value is assigned, if a pair of amino acids does not exist in a sequence.

### Sequence Clustering and Secondary Structure Analysis

Sequence clustering was done at different identity thresholds using MMSeqs2 (Steinegger and Söding, 2017). For sequence homology detection, default MMseq2 (Santoni et al., 2016) settings was used to perform pairwise alignment between all possible pairs of generated and natural sequences in training data. To assess the structural similarity, SPOT-1D (Hanson et al., 2019) was used to predict 3 state (helix, strand, coil) secondary structure for all amino acids. For outlier detection in secondary structure ratio, given the sum of all three ratios is 1, a bivariate Gaussian distribution was fitted only on the helix and strand percentage of natural hexons. Mahalanobis distance (*d*) (Etherington, 2019; Mahalanobis, 1936) was calculated between all generated samples and the center of bi-variate Gaussian distribution.Since *d*^2^follows Chi-square distribution, critical value *α* = 0.05 was used to determine the cutoff distance. Samples with the smallest Mahalanobis distance and the smallest maximum sequence identity are identified to form the Pareto front (Teich, 2001). For selected sequences, their 3D structure was predicted using Alphafold2 (Jumper et al., 2021)

### Sequence Repetitiveness

In order to compare sequence repetition patterns across different positions throughout the sequence, a sliding window with a fixed length was defined. For a sequence of length *L*, there are a total of *L* − *l* + 1 possible positions for the start of window, each position can be normalized *norm*_*pos* = *pos* / *L*. All windows were separated into 20 bins according to the normalized start of their start position. In each window, the number of repeated amino acids was counted. Mean repetition number and mean start of window position were plotted for all 20 bins.*l* was defined. For a sequence of length, there are a total of *L* − *l* + 1 possible positions for the start of window, each position can be normalized *norm*_*pos* = *pos* / *L*. All windows were separated into 20 bins according to the normalized start of their start position. In each window, the number of repeated amino acids was counted. Mean repetition number and mean start of window position were plotted for all 20 bins.

### Latent Space Clustering and Interpolation

To visualize the latent space, Principal Component Analysis (PCA) (Abdi and Williams, 2010) was used to reduce the number of dimensions to 2. 10 biggest clusters (457 sequences in total) at 90% sequence identity in natural hexon sequences were used for plotting.

For interpolation, 1000 points were linearly sampled between the hidden vector of adenovirus 2 hexon and adenovirus 5 hexon. Hidden codes were passed through decoder to get the predicted probabilities. At each position, the token with the highest logit is chosen, and average positional likelihood is calculated as the mean of the max logit across all positions.

As a control, direct interpolation between two sequences was done, by sampling a Bernoulli random variable with *α* probability to choose amino acid from adenovirus 2 hexon (1 − *α* probability from adenovirus 5 hexon) at each position. 1000 different *α* are linearly selected from 0 to 1. 10 sequences were sampled at each *α*, resulting in 10000 sequences in total. ProteinVAE was run on the directly interpolated samples, and collected the predicted probabilities. The average positional likelihood is instead calculated as the mean of the logit for the input sequence. *α* probability to choose amino acid from adenovirus 2 hexon (1 − *α* probability from adenovirus 5 hexon) at each position. 1000 different *α* are linearly selected from 0 to 1. 10 sequences were sampled at each *α*, resulting in 10000 sequences in total. ProteinVAE was run on the directly interpolated samples, and collected the predicted probabilities. The average positional likelihood is instead calculated as the mean of the logit for the input sequence.

### ProtGPT2 Fine-tuning and Sampling

For comparison, ProtGPT2 model was fine-tuned on our training data.Due to GPU memory limitation, 8 AMD MI50-32GB GPUs were in parallel, with a total effective batch size of 16. Learning rate from 10^−6^ to 10^−4^ was tested. The final model used for generation was trained with a learning rate of 10^−5^. The fine-tuned model was prompted with “M” at the start of sentence, and the sampling settings published in the original article was used (Ferruz et al., 2022) for generation. It was observed that generation performance drastically worsened with inclusion of any token with “X”, and all such tokens were removed. The maximum token allowed to place in a sequence is set to 1000 to accommodate the case of the model using single amino acid tokens only. Inference of 50 sequences was repeated for 851 batches (42,550 sequences in total, ∼13h inference time), until 1500 sequences within the length range of hexon were accumulated. Sequences were ranked according to their perplexity (Jelinek et al., 1977), and only kept the top 1000 for comparison. To accommodate downstream analysis, “Z” and “B” found in ProtGPT2-generated sequences were replaced with appropriate standard amino acids.

### ESM-IF1 Generation

100 natural hexons were randomly selected from the training data, and their structures were computationally predicted with Alphafold2 as described above. For each computationally obtained structure template, 100 sequences were generated with ESM-IF. The 10000 generated sequences were ranked according to their log likelihood, and only the top 1000 sequences were used for comparison.

### Molecular Dynamics Simulation Setup

Input structures were used to build the protein-membrane system using CHARMM-GUI online server (Lee et al., 2016). Systems were solvated in an explicit TIP3P (Jorgensen et al., 1983) water box. Charge neutrality was maintained by addition of counter ions, and physiological condition was mimicked using 0.15 M KCl.

All systems were energy minimized using steepest descent before pre-equilibration phase, which was conducted for 500 ps under the constant number of particle, volume, and temperature (NVT) condition. For each system, the production phase was carried out for 100 ns. The particle-mesh Ewald (PME) (Darden et al., 1993; Essmann et al., 1995) method was used with a cut-off radius of 1.2 nm for long-range electrostatic interactions. Heavy atom-hydrogen atom bonds were constrained using the parallel linear constraint solver (P-LINCS) algorithm (Hess, 2008). The Nosé-Hoover thermostat (Hoover, 1985) with a coupling time constant of 1 ps and the Parrinello-Rahman barostat (Parrinello and Rahman, 1981) with a coupling time constant of 5 ps were used for the production phase. A reference coupling pressure of 1 bar and a compressibility of 4.5 × 10^−5^ bar^−1^ were used. For all MD simulations, periodic boundary conditions were applied in all directions. Simulations were carried out using CHARMM36m force field (Huang et al., 2017) by GROMACS/2021.3 (Lindahl et al., 2021). Structure visualization (Figure 4) was done using Protein Imager (Tomasello et al., 2020).

### Human AdV Hexon Classifier

All training sequences were labeled as human or non-human AdV hexon according to their fasta description. All training samples were converted to latent vectors with the previously trained encoder. A simple logistic regression classifier is trained to predict whether a training sequence is from human or non-human AdV from its latent vector. Learning rate is set to 5e^−5^. All samples previously generated from each cluster were encoded, and the trained classifier predicted whether they come from human AdV from their latent vectors.

### Imputed serotyping

Imputed serotyping was conducted as previously described (Madisch et al., 2005). All natural human AdV hexons with known serotypes were extracted and aligned with all generated sequences that are predicted to be human AdV hexon. Location of hexon loop1 and loop2 were identified according to the sequence reported (Madisch et al., 2005) Pairwise amino acid divergence was computed for all possible pairs in the MSA to distinguish if generated sequences are from any known human serotypes. It was reported that amino acid divergence higher than 4.2% in Loop1 and 2.1% in Loop2 to all previously reported serotypes supports that a new serotype is likely identified.

