## Supplemental figures for "ProteinVAE: Variational AutoEncoder for Translational Protein Design"

### Supplementary Figures

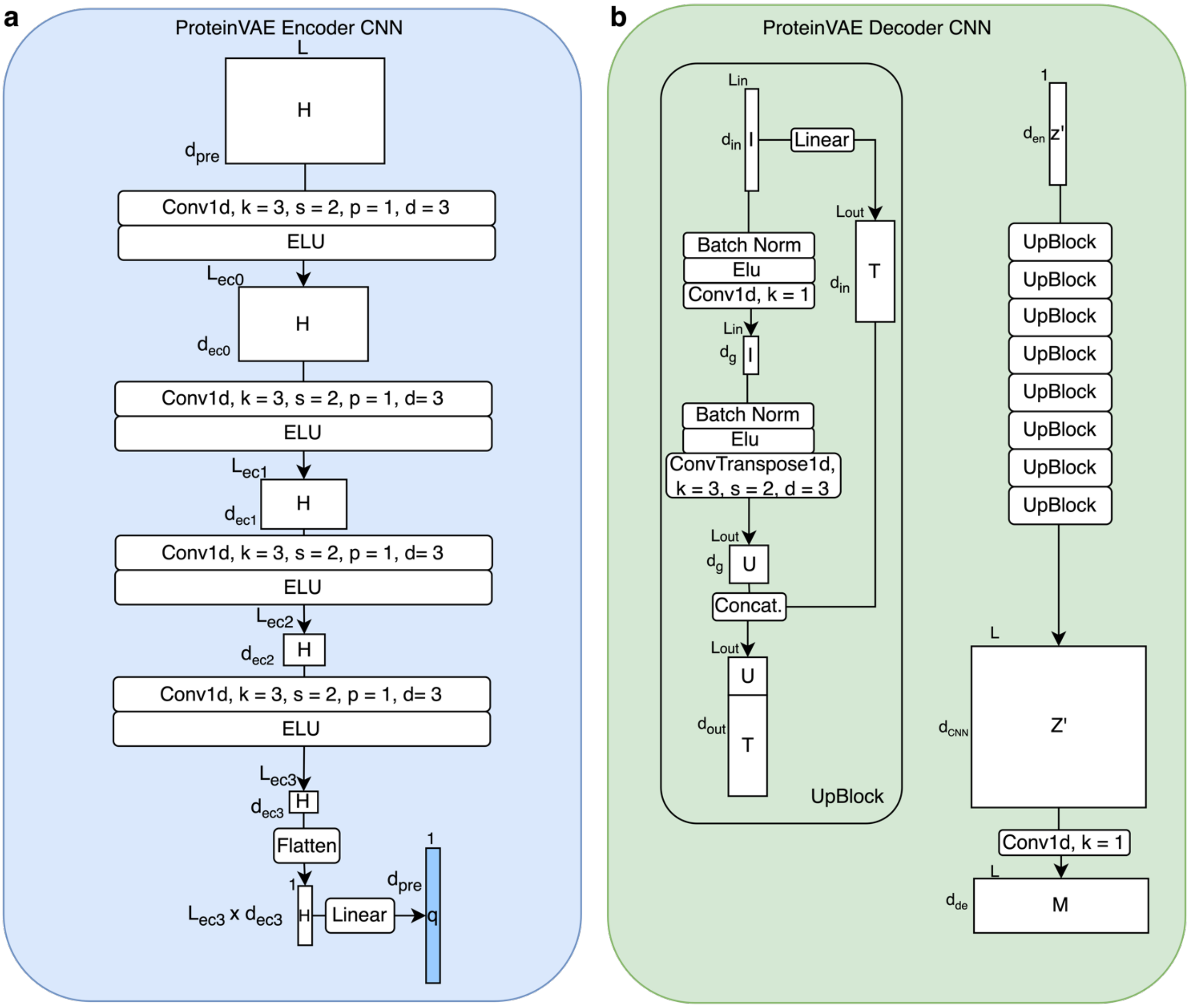


**Supplementary Figure 1 (a)** Encoder CNNs used a series of dilated 3x3 convolution layers along the sequence length dimension to reduce dimensionality of the pretrained language model amino-acid level embeddings. The flattened matrix is then transformed to the same length as the latent size of the pretrained language model embeddings to be used as the query in bottleneck attention. **(b)** Decoder CNNs used 8 UpBlocks to upsample the VAE latent vector length (equals 1) to maximal sequence length. In each UpBlock, a 1x1 convolutional layer is used to transform input to a lower dimension, which reduces the number of parameters needed in the following layer with large kernels. The dilated 3x3 deconvolutional layer with stride of 2 is used to upsample the low-dimensional input. To prevent gradient vanishing, the input is also passed through a linear layer to get an identity matrix (T) of the same length as the deconvolutional output (U). The upsampled matrix U and the identity matrix is then concatenated as the input for the following UpBlock. The output of the final UpBlock is transformed to the decoder hidden dimension with another 1x1 convolutional layer.

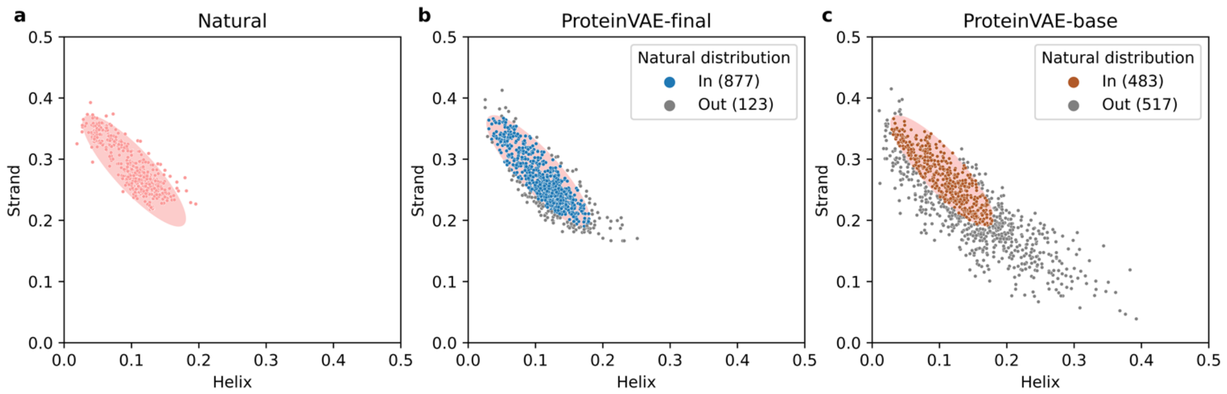


**Supplementary Figure 2** **(a)** Natural hexon helix-strand ratio. **(b)** helix-strand ratio from sequences generated using the final version of ProteinVAE model. **(c)** helix-strand ratio from sequences generated using the base version of ProteinVAE model. Secondary structure state was predicted from sequences directly using SPOT-1D.

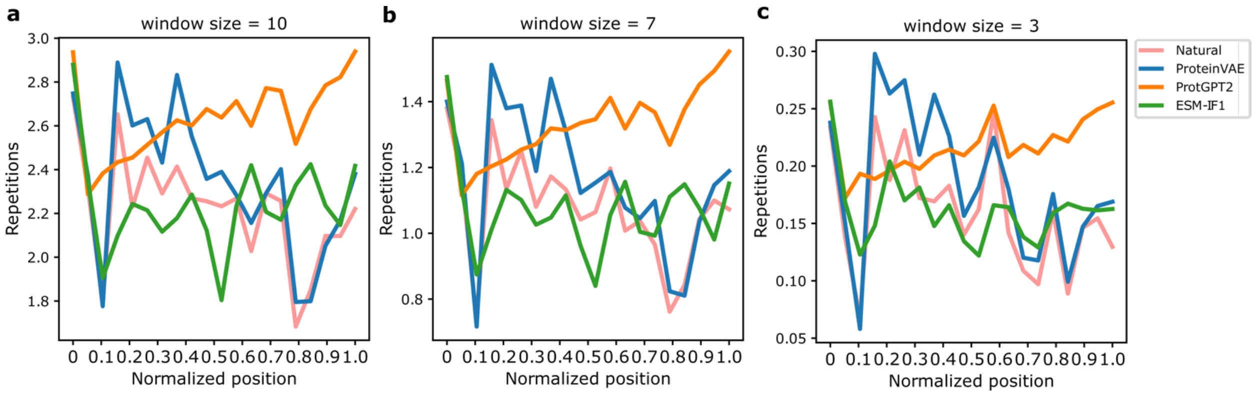


**Supplementary Figure 3** (**a - c**)**:** Repetition profile for natural, ProteinVAE generated, ProtGPT2 generated, and ESM-IF1 generated sequences with window size of 10, 7, and 3 respectively.

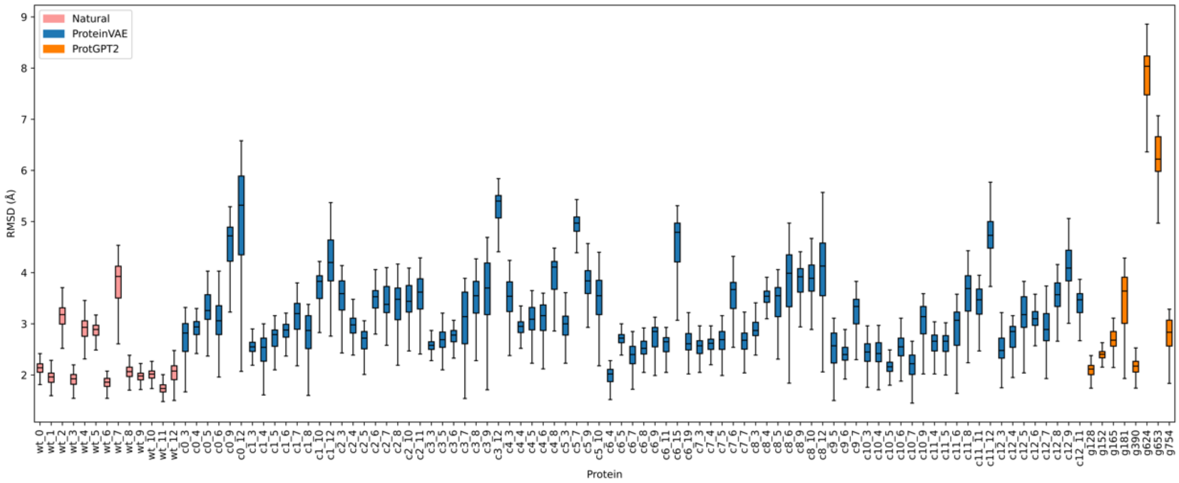


**Supplementary Figure 4** RMSD for all natural representative sequences, ProteinVAE generated sequences, and ProteinGPT2 generated sequences.

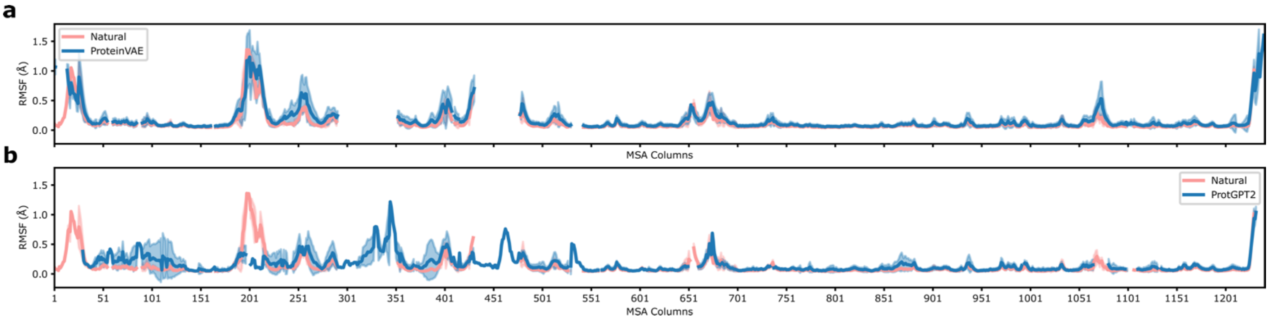


**Supplementary Figure 5 (a)** Average RMSF for ProteinVAE generated sequences (blue) and natural representative sequences (pink) **(b)** Average RMSF for ProtGPT2 generated sequences (blue) and natural representative sequences (pink). ProtGPT2 generated sequences inserted long fragments that are not homologous to any natural hexon. These fragments also have increased flexibility which could reduce structure stability.

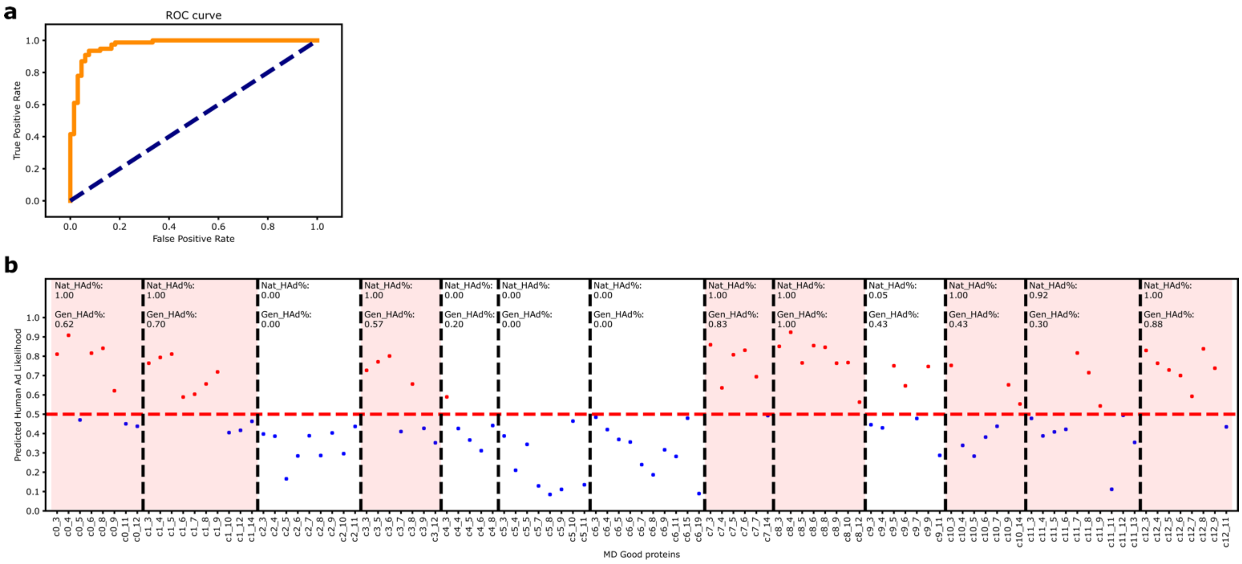


**Supplementary Figure 6 (a)** Receiver operating characteristic (ROC) curve of latent human adenovirus hexon classifier. Area under the ROC curve is 0.97. **(b)** Predicted human AdV hexon likelihood for all sequences generated from each cluster. Sequences predicted to be human AdV hexon were shown as a red dot, and predicted non-human AdV hexon were shown as a blue dot. Percentages of human AdV in corresponding natural sequences were labeled as Nat_HAd% in each cluster. Clusters with more than 90% natural human AdV hexons were colored with a pink background. Predicted percentages of human AdV for generated sequences were labeled as Gen_HAd%. Decision threshold is shown as a dashed red line.

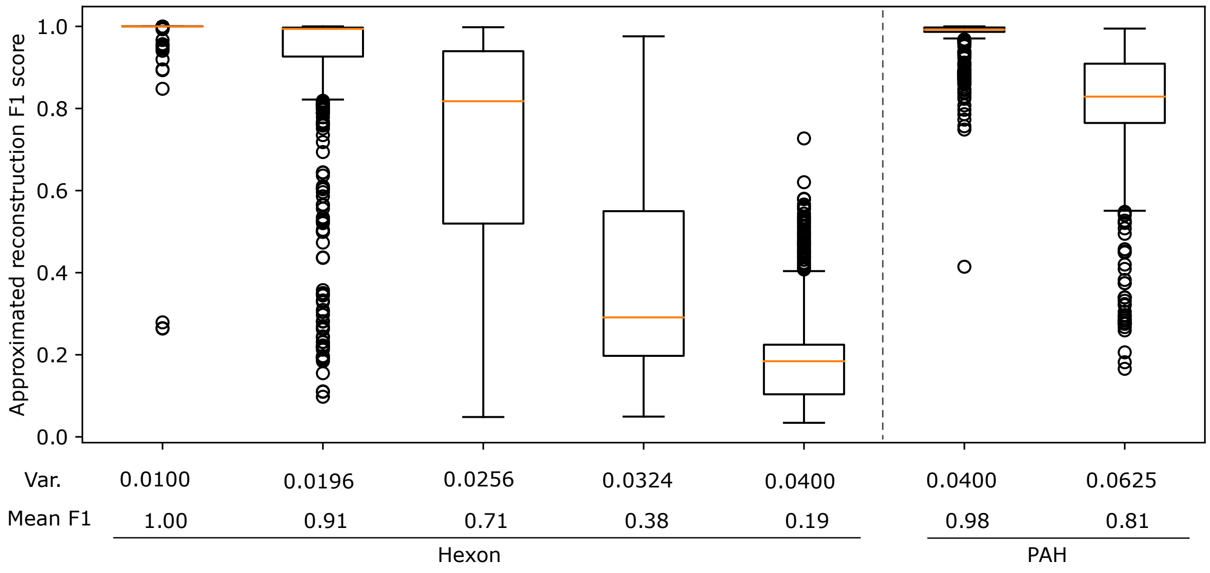


**Supplementary Figure 7** Approximated reconstruction F1 score is compared between hexon dataset and small PAH dataset extracted the same way as described in ProT-VAE. Small Gaussian noises with different variance were introduced to the encoder output to simulate inaccuracy injected by the CNNs and VAE component in ProT-VAE. As can be seen, the simulated reconstruction F1 worsened even at very low variance for the hexon dataset, but the reconstruction performance was maintained even at higher variance in the PAH dataset.
